# Probabilistic Modelling of Base Editing Efficiency and Bystander Outcome Composition

**DOI:** 10.64898/2026.10.09.757916

**Authors:** Furkan Özden, Peter Minary

**Affiliations:** Department of Computer Science, University of Oxford, Parks Rd, Oxford OX1 3QD, United Kingdom

## Abstract

Base editors install single-nucleotide substitutions without a double-strand break or donor template, but editing at multiple susceptible positions can generate bystander alleles. Predicting the frequency of an intended outcome therefore requires both editing efficiency and the composition of edited products. Here we present crispAIBE, a probabilistic framework that uses separately trained transformer– convolutional networks to predict Dirichlet distributions for editing efficiency and conditional allele composition. Split-conformal density-level regions are calibrated on data reserved from model selection, with composition thresholds fitted by outcome-set size. On held-out data, crispAIBE achieves Pearson correlations of 0.669 for efficiency and 0.896 for composition on measured allele sets, with empirical coverages of 0.902 and 0.906 at a nominal level of 0.90. On the published CRISPRon-BE benchmark, with test protospacers excluded from training, crispAIBE ranks first in seven of eight comparisons under the two-column Pearson statistic and first or second in seven of eight under the rank statistic. Within predicted-efficiency quintiles, the largest- and smallest-region deciles differ 1.9-fold in mean absolute error on average, compared with 1.7-fold for a boundary-distance ranking. When evaluated on U2OS, which was excluded from training, larger predictive regions were associated with larger absolute prediction errors. Local calibration with 100 U2OS protospacers yields median empirical coverage of 0.918 at the nominal 0.90 level. The trained crispAIBE models and scripts for data processing, training, calibration and evaluation are available at https://github.com/furkanozdenn/crispAIBE.

## 1 Introduction

Base editors install single-nucleotide substitutions without a double-strand break or a donor template [3]. Cytosine base editors (CBEs) typically convert C· G to T· A [13], whereas adenine base editors (ABEs) convert A· T to G· C [6]. Their therapeutic potential has been demonstrated by a personalised in-vivo treatment of an infant with a urea-cycle disorder [20]. A guide may expose several susceptible bases within the editor’s activity window. The resulting alleles differ in which positions are converted. Guides with similar overall efficiency can therefore produce different proportions of the intended edit and bystander products. The fraction carrying a particular edited allele depends on both the overall edited fraction and that allele’s proportion among edited products. Estimating these quantities separately distinguishes low editing activity from efficient editing that favours unwanted products.

The distinction also requires information about which edits occur together. Per-position editing frequencies alone do not generally determine allele composition: the same marginal frequencies can arise from different combinations of edits. Representing these combinations explicitly makes the joint pattern of edited positions a prediction target alongside overall efficiency. This matters when an intended conversion can occur either alone or together with bystander changes.

BE-dataHIVE compiles editing-efficiency and bystander data from multiple studies in standardised formats for model development [27]. Existing predictors address efficiency, outcome composition, or both. BE-Hive combined large target-library screens with models of guide efficiency and allele frequency [4]. Subsequent approaches include attention-based BE-DICT [18], per-position prediction with FORECasT-BE [23], endogenous-context modelling with BE_Endo [35], and prediction across larger editor panels with DeepBE [11]. BEDICT2.0 evaluated prediction across cell lines and mouse liver [12]; BEguider addressed near-PAMless editors [36]; and BEEP compared high-activity adenine and cytosine editors [34]. CRISPRon-BE combined several screens using dataset-origin labels and reported a benchmark against existing predictors [30]. These methods provide estimates of editing outcomes, but do not establish calibrated guide-level predictive regions for both efficiency and conditional bystander composition.

Predictive uncertainty complements point estimates by describing the dispersion of possible outcomes before experimental validation [5,10]. Within genome editing, uncertainty has been studied for Cas9 off-target activity [22] and cut efficiency [26]. Our earlier crispAIPE model combined Dirichlet likelihoods and conformal regions for a fixed composition of intended edits, unedited reads and pooled unintended products [21]. Prime-editing predictors such as PRIDICT2.0 and PrimeNet also estimate outcome proportions [19,16].

Base-editing composition introduces guide-dependent combinations of edited positions, so the identities and number of represented outcomes vary across guides. The composition network in crispAIBE scores each represented allele pattern and assembles its concentrations into a guide-specific Dirichlet distribution. This construction accommodates variable-size allele sets while modelling their proportions jointly. Calibration by outcome-set size accounts for the different density scales on the corresponding simplices.

Guides with similar predicted means can have different predictive dispersions. Predictive regions describe this dispersion while respecting the constraint that allele proportions sum to one. We assess their calibration by empirical coverage and their informativeness by the association between region size and prediction error.

We developed crispAIBE to model efficiency on the edited/unedited simplex and composition on a conditional simplex over allele patterns. Each model produces a Dirichlet distribution, whose concentration describes the dispersion of its predictions [28,17]. We then calibrate density-level regions using held-out data [8,32,24,14]. Our evaluation separates three questions: prediction accuracy, empirical region coverage, and whether region size is associated with prediction error. We also compare results on the benchmark published by the CRISPRon-BE authors, examine the contributions to its combined metrics, and test uncertainty when U2OS cells are excluded from training.

## 2. Materials and Methods

### 2.1 Datasets and splitting

We assembled 524,508 records associated with 457,017 source-specific guide identifiers and 454,980 distinct protospacer–editor–cell combinations. The sources were BE-Hive’s per-guide efficiency and per-allele genotype releases [4] (figshare articles 10673816 and 10678097), and the compilation accompanying CRISPRon-BE [30], which includes SURRO-seq, Song, Marquart and Kissling data. We used these releases to retain the allele-level information required for composition modelling. The corpus contains 17 editor labels across HEK293T, U2OS and mES cells. It includes 127,533 ABE records and 396,975 CBE records, of which 10,607 derive from the transversion-enriched CtoGA library. The latter is a library category, not a separate molecular editor class.

The efficiency target is the observed edited fraction. Where genotype measurements are available, the composition target is the distribution of allele patterns conditional on editing. The main composition dataset excludes guide-context observations with fewer than 100 edited reads; the multi-source benchmark composition dataset uses a threshold of 50. The main corpus uses approximately 88%, 5% and 7% train, validation and test partitions, with all records of a protospacer assigned together; Marquart records are assigned to test.

Before removing missing efficiency labels, these partitions contain 463,709, 25,781 and 35,018 records. The fitted efficiency model uses 463,517 labelled training records associated with 403,006 guide identifiers. Its test metrics use 35,006 labelled records. The main composition model trains on 216,211 guide-context observations containing 1,215,821 allele records. We distinguish these observations from unique protospacers, which may occur in several editing contexts. The U2OS cell-line transfer experiment uses a different split: training and validation retain their original assignments after excluding U2OS, whereas evaluation uses all U2OS records. This experiment can include protospacers observed in training under another cell context (Section 3.8).

*External benchmark exclusion*. We excluded every protospacer in CRISPRon-BE’s partition-6 test files from both benchmark models’ training data. For efficiency, this excludes the benchmark’s 7,912 test protospacers. For composition, exclusion also covers BE-Hive, which shares 3,077 protospacers with that partition; removing these eliminates 53,405 BE-Hive guide-context rows. The benchmark composition model uses 222,290 training guide-context observations. The training corpus and the external partition are checked for exact protospacer overlap. Marquart is absent from the CRISPRon-BE training files and is an external source for both methods. Arbab is omitted from comparative claims because it is the same underlying screen as BE-Hive.

The benchmark composition checkpoint was selected using a multi-source validation split that retained source-specific identifiers. Unlike the main composition split, its internal partitions are not fully protospacer-disjoint: 2,553 sequences occur in both training and validation, and 322 in training and its internal test split.

We therefore use this checkpoint only for the separately excluded external benchmark, and do not interpret its internal validation or test performance as generalisation to unseen protospacers.

### 2.2 Modelling

#### Problem formulation

Let a guide be a 20-nt protospacer *s* edited by editor *e* in cell context *c*. We predict two compositional quantities. The first is the efficiency composition **y** = (*y*_ed_, 1 −*y*_ed_) on the 1-simplex. The second is the conditional bystander composition ***π*** = (*π*_1_, …, *π*_*K*_ *s*) on the (*K*_*s*_ − 1)-simplex, where the allele set of size *K*_*s*_ contains the represented edit patterns and, where applicable, an aggregated remainder. We model both as compositions rather than as independent scalars. For each target, a Dirichlet likelihood enforces the constraint that outcome proportions sum to one, and calibration defines a region on the corresponding simplex.

#### Allele representation

We restrict the represented activity window to protospacer positions 3–10. Each allele pattern is an eight-bit vector indicating which positions are edited, together with a ninth component marking a residual category. The main dataset aggregates alleles by their in-window pattern, collapsing distinctions outside the window. It retains the seven most abundant measured patterns separately and combines the remaining mass in an “other” slot, encoded by eight zero bits and a residual flag. There are therefore at most eight valid slots per guide. Edits at the same position are collapsed to a single bit, so this representation does not distinguish substitution identities or describe indels explicitly. In the BE-Hive data, every reported in-window edit sets its positional bit, including an edit at a non-target base. The ninth component in the main dataset marks only the aggregated remainder, rather than a particular class of molecular change.

The benchmark representation also flags changes outside the represented window. For its CRISPRon-BE records, edits to the editor’s target base within positions 3–10 set positional bits; an edit outside the window or at a non-target reference base sets the residual flag. For example, a CBE substitution of an in-window cytosine to T, A or G sets the same positional bit: a noncanonical substitution does not itself activate the ninth component. Such products have the same positional encoding as the canonical edit and are not distinguished by substitution identity. For BE-Hive records in the multi-source dataset, every in-window edit sets a positional bit and changes outside the window set the residual flag. Positional bits and the flag can both be active when an allele contains both kinds of change. An aggregated tail uses the same all-zero positional vector and residual flag as in the main dataset. Thus the ninth component can indicate an unrepresented change or aggregated mass, depending on the record.

For a canonical-only illustration, consider a guide with *m* = 4 editable target bases and 2^4^ − 1 = 15 possible non-empty patterns. If all fifteen are observed, the seven most abundant are selected across all four positions; the selection does not restrict editing to three positions. The other eight are aggregated into “other”, encoded as 000000001 although all are canonical in-window edits. This choice limits the number of separately modelled components while preserving the total measured mass. When at most seven patterns are observed, all are retained individually and no remainder slot is needed.

The top-*K* selection uses measured abundance in training, validation and test. The reported main composition accuracy and coverage therefore concern retrospective prediction on measured allele sets. For a new, unmeasured guide, the observed top seven are unknown. Sequence enumeration supplies 2^*m*^− 1 non-empty candidate patterns to the per-allele scorer for a guide with *m* editable target bases. This changes the normalisation and potentially the dimension of the predictive distribution. The reported top-*K* results are specific to the measured allele representation; they do not quantify accuracy or coverage for sequence-enumerated candidates or prediction-based top-seven selection. The external benchmark is evaluated separately on the outcome sets supplied in its released files.

The model receives allele identity but no slot index. Permuting allele slots therefore permutes the predicted concentrations. We use this property to assess the contribution of packing order to the reported composition correlations.

#### Architecture

Efficiency and composition are predicted by two separately trained networks with related architectures; they do not share fitted encoder weights. Each network embeds the 20-nt protospacer, adds sinusoidal positional encodings, and applies a two-layer transformer with four attention heads [31]. Transformer outputs are concatenated with the one-hot sequence and passed through a convolutional block, followed by max-pooling over positions. The efficiency network additionally supplies protospacer, activity-window and editable-base masks to the convolutional block. Editor and cell embeddings of dimensions 16 and 8, respectively, are concatenated with the pooled representation. The main efficiency and composition models use vocabularies of 17 and 13 editors, respectively. The main models use no auxiliary scalar features. The benchmark efficiency model and the U2OS transfer model additionally receive GC content, predicted guide–target hybridisation energy and activity-window GC content. In those experiments, feature means and standard deviations are calculated over the full filtered input corpus before splitting, including test inputs. Editor and cell vocabularies are also defined over the full corpus. This preprocessing uses input features and category names, without test outcome labels.

The efficiency output is a positive two-component concentration vector, produced by a feed-forward network with softplus outputs. Its Dirichlet distribution is equivalent to a Beta distribution over the edited fraction. The composition network treats allele identity as an input rather than a fixed output-class label. For each guide, its context vector is concatenated with each nine-component allele pattern and passed through the same feed-forward scorer, producing one positive concentration per allele. These concentrations parameterise a single Dirichlet distribution over the supplied allele set; a mask excludes unused padding from the likelihood. The same scorer therefore accommodates guide-specific allele sets of different sizes, with parameters shared across alleles within the composition network.

For either model, normalising the concentrations over valid components gives the predicted mean proportions. In particular, the composition mean is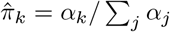, where the sum runs over valid alleles. For distributions with at least two components, scaling all concentrations together leaves these means unchanged but changes the dispersion: at fixed means, larger total concentration gives smaller componentwise variance. This separates the predicted composition from the model’s dispersion around it. For either target, the negative log-likelihood is

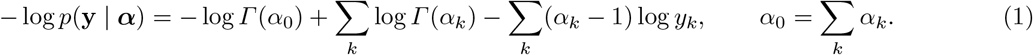

These are fitted conditional predictive distributions, not Bayesian posteriors over network parameters. Efficiency targets are clipped to [10^*−*4^, 1 − 10^*−*4^] by the data loader; composition proportions are clipped at 10^*−*6^ when evaluating logarithms in the loss. Both networks use Adam with learning rate 6 × 10*−*^4^, batch size 256 and early stopping on validation loss. The main networks use no weight decay.

Because BE-Hive contributes most records, efficiency training for the external benchmark uses source-balanced sampling, with weight proportional to 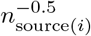for record *i*. In a separate benchmark training trial, full inverse-frequency weighting attained its best validation loss after the first epoch. The main in-distribution and ablation experiments, and the U2OS transfer experiment, use unbalanced sampling.

#### Conformal region construction

A model-based highest-density region contains the specified probability mass under the predicted Dirichlet distribution [8]. Its coverage of observed outcomes therefore depends on the predictive distribution. In the calibrated construction, the Dirichlet density determines the ordering of possible compositions, while held-out observations determine the density threshold used to form the region. We compare these model-based and calibrated constructions on the same fitted predictor.

For a fixed predictor, let *S*_*i*_ = − log *p*(**y**_*i*_ | ***α***_*i*_) for *n* calibration observations and let *k* = ⌈(*n* + 1)*γ*⌉ for target coverage 0 *< γ* ≤ 1. The implemented threshold is

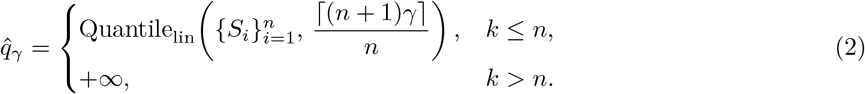

Here Quantile_lin_ uses linear interpolation between ordered scores at zero-based index (*n* − 1)*k/n*. The standard split-conformal threshold is the *k*th ordered calibration score [32,15,14]; for *k ≤ n*, the interpolated threshold in Eq. 2 is at least as large. The resulting density-level set is

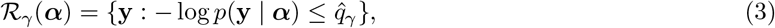

This raw-density score differs from the integrated density-rank score used by HPD-split [9].

For composition, raw density scores have different scales on simplices of different dimensions. We therefore fit a separate threshold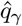_,*K*_ for each represented set size *K >* 1. In Eq. 2, we replace *n* by the count *n*_*K*_ of calibration records with set size *K*, and use only their scores {*S*_*i*_ : *K*_*i*_ = *K*}. For *K* = 1, the region consists of the sole possible composition (1). If a set size is absent from calibration, or its required quantile rank exceeds the available count, the region is the entire simplex. Efficiency has fixed dimension and uses a pooled threshold unless editor stratification is specified.

For exchangeable calibration observations and a future observation, with the predictor fixed independently of calibration, the construction provides marginal coverage at least *γ* [29,2]. Our corpus contains repeated protospacers across editors, cells and measurements. We keep a protospacer’s records together when splitting the main test data, but calculate calibration quantiles and coverage over records. For this clustered design, the observation-level theorem does not directly imply finite-sample record-level coverage. We therefore report empirical coverage for the specified evaluation population.

For the main coverage analysis, a deterministic hash of the protospacer partitions the test set into calibration and evaluation halves, neither used for model selection. Efficiency calibration uses 16,935 records from 2,637 protospacers and evaluation uses 18,071 records from 2,800 protospacers. The corresponding composition counts are 7,755 records from 1,203 protospacers and 8,267 from 1,218. Accuracy uses the full test split. The validation-fold calibration protocol reuses observations involved in early stopping. The head and backbone ablations and uncertainty-ranking analyses retain that protocol. The region-construction comparison uses the disjoint test halves.

Region size is the fraction of the simplex enclosed, estimated by uniform Monte Carlo sampling. For the efficiency model this is the length of the region within [0, 1]. We also evaluate editor-stratified (Mondrian) calibration, fitting a separate threshold within each editor [33]; its theoretical within-editor guarantee requires exchangeability within those strata.

### 2.3 Evaluation

For the benchmark comparison, we used the two-dimensional Pearson and Spearman rank correlation coefficients, *R*_2_ and *ρ*_2_, retaining the terminology of Sun et al. [30]. These statistics derive from the *K*-category correlation coefficient *R*_*K*_ of Gorodkin [7], which generalises the Matthews correlation coefficient (equivalently Pearson’s *ϕ*) from two categories to *K*. Gorodkin defines *R*_*K*_ for a pair of *N*× *K* matrices, of which the familiar confusion-matrix statistic is the special case where those matrices hold categorical assignments; Sun et al. evaluate the same covariance form on continuous columns at *K* = 2, taking the two columns to be gRNA editing efficiency and outcome frequency, which gives *R*_2_. The rank form *ρ*_2_ is introduced in the same work, and reduces to the Spearman coefficient at *K* = 1. Each (guide × outcome) pair contributes an editing-efficiency column and an outcome-frequency column. Section 3.4 describes the variance-dependent coefficients and how ranking changes them. The coefficient is

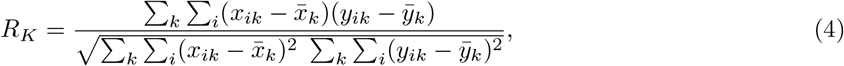

evaluated on the raw values (*R*_2_) or on within-column ranks (*ρ*_2_). Predicted outcome frequencies are formed by combining the separately trained models’ predicted means: a wild-type row receives 100(1 − Ê) and an edited allele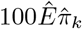, where Ê is the predicted efficiency. Ground truth is CRISPRon-BE’s published per-outcome frequency, and we compare against their best-performing dataset-label variant per test set as reported in their Supplementary Data 3. The best variant is selected separately for each reported metric.

## 3 Results

### 3.1 Overview of crispAIBE

crispAIBE comprises an efficiency model and a conditional composition model (Fig. 1). Their inputs identify the protospacer, editor and cell context; the composition model additionally receives each candidate allele pattern. The efficiency model predicts a distribution over edited fraction, and the composition model predicts a distribution over the proportions of the represented patterns. Separate calibration data set the density threshold used for each predictive region.

**Fig. 1.**
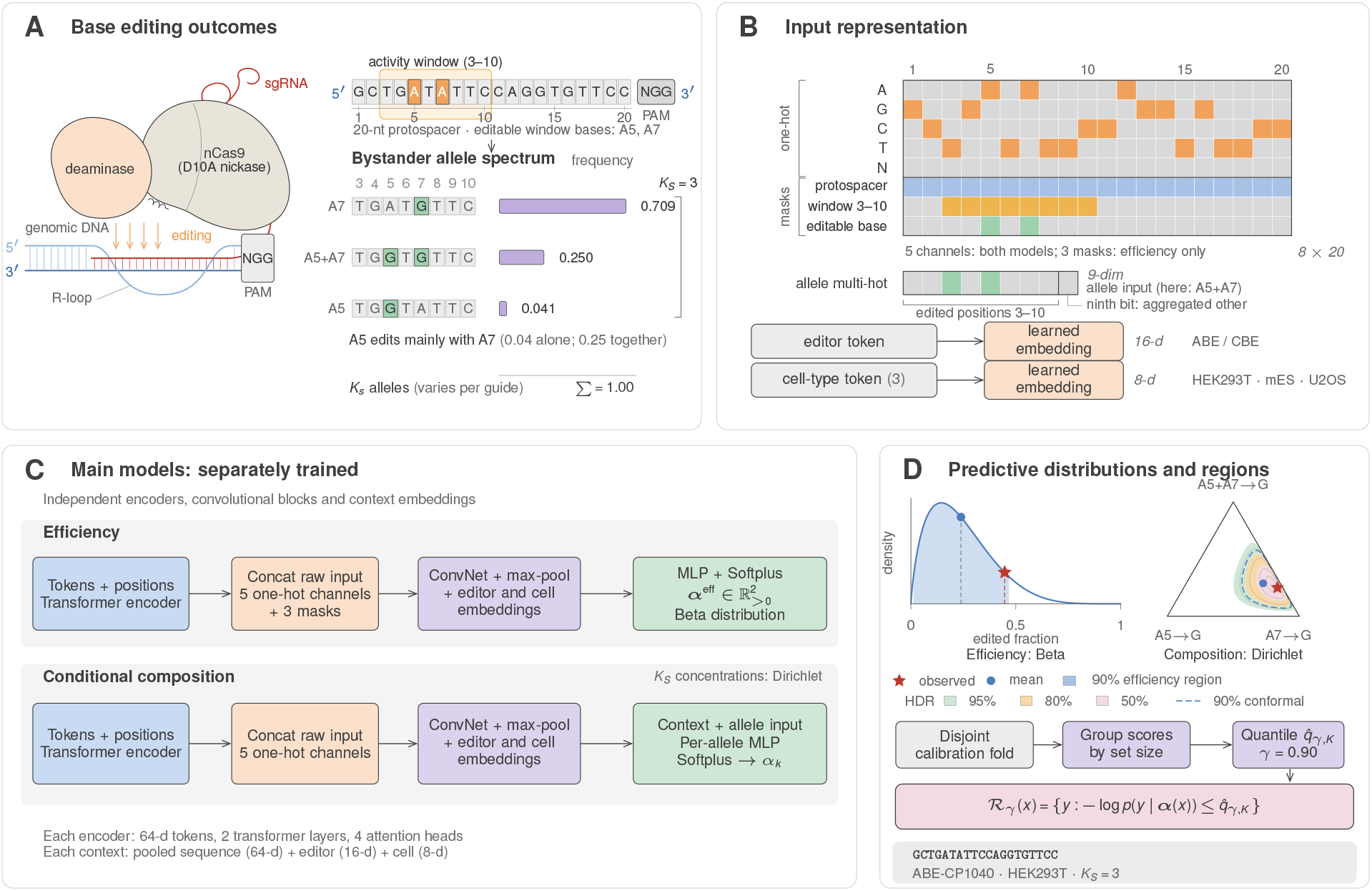
crispAIBE models for efficiency and conditional composition. The two networks are trained separately with related transformer–convolutional architectures. They predict Dirichlet distributions on the edited/unedited simplex and the simplex of represented bystander patterns, respectively. Panel **C** depicts the main models. Calibration thresholds are fitted on the disjoint test-half fold described in Methods; *K* denotes the number of components, fixed at two for efficiency and equal to *K*_*s*_ for composition. Panels **A, B** and **D** follow one held-out test guide throughout (protospacer GCTGATATTCCAGGTGTTCC, ABE-CP1040, HEK293T), whose activity window contains two editable adenines at protospacer positions 5 and 7 and whose observed allele set has cardinality *K*_*s*_ = 3. Its A5 edit occurs mainly together with A7 (0.04 alone against 0.25 jointly). The predictive distributions drawn in **D** are the model’s actual predictions for that guide.

The efficiency corpus comprises 494,166 BE-Hive records and 30,342 records from the CRISPRon-BE compilation. The main composition dataset contains 1,374,730 allele records over 244,015 guide-context observations. The benchmark composition dataset contains 1,541,730 allele records over 247,486 guide-context observations after external-test exclusion. Editor names are canonicalised across sources.

### 3.2 Accuracy and empirical coverage on held-out records

The efficiency model attains Pearson *r* = 0.669 and Spearman *ρ* = 0.677 on the full held-out test split. Empirical coverage on the separate evaluation half is 0.902 at the nominal 0.90 level (Fig. 2A–B). For composition, per-allele Pearson correlation is 0.896 and Spearman correlation is 0.800, with empirical coverage 0.906 (Fig. 2C–D). Composition accuracy and coverage were evaluated on each guide’s measured top-seven patterns and aggregated remainder; all represented entries in this test split had positive observed frequency.

**Fig. 2.**
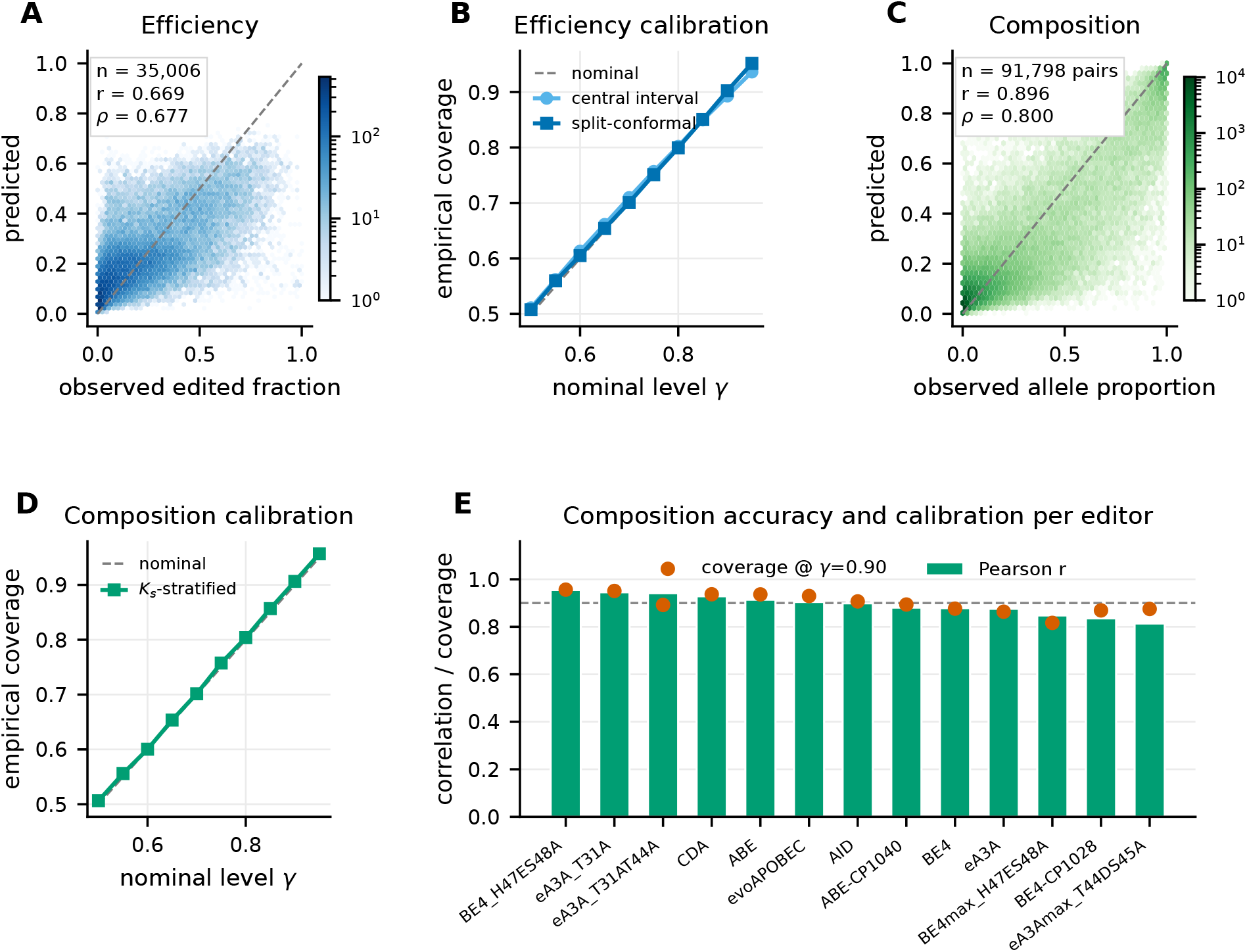
Prediction accuracy and calibration diagnostics for crispAIBE. **(A)** Predicted versus observed edited fraction on the held-out test split (*n* = 35,006 guide-context records), as a log-density hexbin; the dashed line is the identity. **(B)** Empirical coverage against nominal level *γ* for the central predictive interval and split-conformal region, both evaluated on the same disjoint evaluation half. **(C)** Predicted versus observed per-allele composition proportions (91,798 guide *×* allele pairs), using the measured top-seven patterns and aggregated remainder. **(D)** Composition coverage against nominal level, using the Dirichlet negative-log-likelihood score calibrated separately by represented set size. **(E)** Per-editor composition Pearson correlation (bars) with conformal coverage at *γ* = 0.90 (points); the dashed line marks the nominal level. Accuracy uses all test records; coverage uses the disjoint evaluation half.

The packing convention creates a strong abundance profile across slots: observed proportions are non-increasing for 80.4% of the 15,803 guide-context observations with at least two represented alleles. For a descriptive reference, we fitted the mean proportion within each (slot, set-size) group directly on the test observations, giving pooled Pearson 0.938. Random slot permutations leave the model’s correlation unchanged at 0.896, while the fitted slot diagnostic falls to 0.329. Among the 14,551 observations with at least three represented alleles, mean within-guide Spearman is 0.735 for the model, compared with 0.969 before and 0.014 after permutation for the slot diagnostic. Removing the respective mean (slot, set-size) profiles from predictions and observations gives residual Pearson 0.596. Thus pooled correlation reflects both an abundance profile shared across guides and variation within that profile. The permutation test preserves the mapping between allele identity and observed proportion while changing storage order. The unchanged model correlation is consistent with its per-allele scoring architecture. The residual correlation measures association after removing the average abundance profile within each slot and set-size group.

Across 13 editors with sufficient test support, composition Pearson correlations range from 0.814 to 0.955 (Fig. 2E). Values are highest for BE4_H47ES48A (0.955), eA3A_T31A (0.945) and eA3A_T31AT44A (0.941), and lowest for eA3Amax_T44DS45A (0.814, *n* = 61) and BE4-CP1028 (0.835). By cell line, Pearson correlation is 0.912 for mES, 0.886 for HEK293T and 0.839 for U2OS. Composition coverage after calibration by set size ranges from 0.816 to 0.956 across editors. Calibration by set size does not impose an editor-specific coverage target.

Pooling raw-density scores across set sizes concealed large differences in coverage. At nominal 0.90, a single threshold pooled across sizes gives coverage 0.900 overall, but 0 for singleton sets and 0.523 for two-component sets. Separate thresholds give coverage 1 for singleton sets and 0.895–0.911 across sizes two to eight (Supplementary Table S3). Overall coverage averages across represented set sizes and can therefore conceal under-coverage within a particular dimension. Calibration by set size compares density scores on the same simplex and resolves this disparity in the evaluated data. Singleton sets require no estimated boundary because their simplex contains only one composition.

### 3.3 External benchmark comparison

We evaluated crispAIBE on the partition-6 test data released by Sun et al. [30], using their two-column correlation statistics. Separate benchmark checkpoints exclude all 7,912 test protospacers from both training corpora, including sequences shared with BE-Hive (Section 2.1). Predictions were available for 79,598 of 79,600 released rows; the two omitted Arbab rows lacked a matching cell-context prediction. Arbab rows are excluded from comparative claims because Arbab and BE-Hive derive from the same screen. The remaining eight source–editor comparisons contain 62,740 outcome rows. Marquart is absent from both our training data and CRISPRon-BE’s released training files. Other sources, including Kissling, contribute separate training guides, so their test sets do not represent unseen laboratories.

crispAIBE has the highest *R*_2_ in seven of eight comparisons and ranks second on Kissling ABE8e (Table 2). Under *ρ*_2_, it ranks first or second in seven of eight comparisons; the mean ranks are 1.12 and 1.75, respectively. Table 1 gives 95% percentile intervals for crispAIBE’s estimates from 2,000 guide-level bootstrap resamples, keeping all outcomes from each sampled guide together. Rankings compare point estimates; paired comparator predictions were unavailable for estimating uncertainty in between-method differences.

**Table 1.** Benchmark point estimates and uncertainty for crispAIBE. CRISPRon-BE values are the highest published dataset-label variant per set and metric [30]. Brackets give crispAIBE’s 95% guide-level bootstrap intervals (*B* = 2000); they are not intervals for the difference between methods. *n* counts guides within each test set. Arbab is excluded because it derives from the BE-Hive screen.

| Editor | Test set | $n$ | $R_2$ | | $\rho_2$ | |
| --- | --- | --- | --- | --- | --- | --- |
|  |  |  | crispAIBE 95% CI | CRISPRon | crispAIBE 95% CI | CRISPRon |
| ABE | SURRO-seq | 2,326 | 0.905 [0.896,0.913] | 0.762 | 0.784 [0.768,0.799] | 0.774 |
| ABE | Song | 430 | 0.887 [0.870,0.904] | 0.804 | 0.825 [0.789,0.853] | 0.847 |
| ABE | Marquart<sup>†</sup> | 1,482 | 0.661 [0.637,0.684] | 0.384 | 0.481 [0.444,0.520] | 0.448 |
| ABE | Kissling ABE7.10 | 144 | 0.759 [0.687,0.831] | 0.715 | 0.572 [0.428,0.700] | 0.620 |
| ABE | Kissling ABE8e | 152 | 0.636 [0.544,0.720] | 0.742 | 0.400 [0.236,0.560] | 0.415 |
| CBE | SURRO-seq | 2,165 | 0.847 [0.833,0.860] | 0.800 | 0.809 [0.793,0.823] | 0.804 |
| CBE | Song | 474 | 0.969 [0.963,0.975] | 0.824 | 0.762 [0.707,0.802] | 0.832 |
| CBE | Marquart<sup>†</sup> | 1,489 | 0.845 [0.825,0.865] | 0.191 | 0.359 [0.322,0.397] | 0.228 |

<sup>†</sup> Absent from both our training data and CRISPRon-BE’s released training files. Kissling ABE7.10 is the published benchmark label; the compiled metadata map these records to ABEmax.

**Table 2.** crispAIBE against every method scored on these test sets. Competitor values are taken from Supplementary Data 3 of Sun et al. [30], who evaluated all of them on the same partition-6 test guides with the same statistic; for CRISPRon-BE we take its best dataset-label variant per test set. A dash means the method was not scored on that set. Bold marks the best value in each row. crispAIBE ranks first on seven of the eight comparisons under *R*_2_ and second on the eighth; under *ρ*_2_ it is first or second on seven of eight. Mean ranks are 1.12 and 1.75 among the methods scored on each set.

|  | crispAIBE | CRISPRon | DeepABE | DeepCBE | DeepBE | BE-Hive | BEDICT2 | BE-DICT | BE_Endo |
| --- | --- | --- | --- | --- | --- | --- | --- | --- | --- |
| $R_2$ (Pearson form) | | | | | | | | | |
| ABE SURRO-seq | **0.905** | 0.762 | 0.665 | — | — | 0.509 | 0.495 | 0.399 | 0.472 |
| ABE Song | **0.887** | 0.804 | 0.721 | — | — | 0.631 | 0.569 | 0.373 | 0.411 |
| ABE Marquart<sup>†</sup> | **0.661** | 0.384 | 0.357 | — | — | 0.354 | 0.344 | 0.297 | 0.221 |
| ABE Kissling 7.10 | **0.759** | 0.715 | 0.441 | — | — | 0.651 | — | 0.489 | 0.237 |
| ABE Kissling 8e | 0.636 | **0.742** | — | — | 0.450 | — | — | 0.151 | — |
| CBE SURRO-seq | **0.847** | 0.800 | — | 0.737 | — | 0.524 | — | 0.304 | 0.494 |
| CBE Song | **0.969** | 0.824 | — | 0.785 | — | 0.586 | — | 0.309 | 0.489 |
| CBE Marquart<sup>†</sup> | **0.845** | 0.191 | — | 0.208 | — | 0.219 | — | 0.211 | 0.146 |
| $\rho_2$ (rank form) | | | | | | | | | |
| ABE SURRO-seq | **0.784** | 0.774 | 0.699 | — | — | 0.590 | 0.546 | 0.476 | 0.563 |
| ABE Song | 0.825 | **0.847** | 0.799 | — | — | 0.728 | 0.662 | 0.529 | 0.592 |
| ABE Marquart<sup>†</sup> | **0.481** | 0.448 | 0.427 | — | — | 0.390 | 0.397 | 0.403 | 0.363 |
| ABE Kissling 7.10 | 0.572 | **0.620** | 0.516 | — | — | 0.513 | — | 0.506 | 0.371 |
| ABE Kissling 8e | 0.400 | **0.415** | — | — | 0.207 | — | — | 0.091 | — |
| CBE SURRO-seq | **0.809** | 0.804 | — | 0.753 | — | 0.585 | — | 0.447 | 0.541 |
| CBE Song | 0.762 | **0.832** | — | 0.811 | — | 0.690 | — | 0.482 | 0.560 |
| CBE Marquart<sup>†</sup> | 0.359 | 0.228 | — | 0.259 | — | 0.198 | — | **0.375** | 0.222 |

<sup>†</sup> Absent from CRISPRon-BE’s training files, which we verified directly, and reported by Sun et al. as external for the remaining methods.

The rankings differ between the Pearson and rank forms (Fig. 3A–B). For example, on CBE Song crispAIBE has *R*_2_ = 0.969 against CRISPRon-BE’s 0.824, but *ρ*_2_ = 0.762 against 0.832. On Marquart, crispAIBE has higher point estimates under both statistics for both editor classes. For Marquart ABE, *R*_2_ is 0.661 for crispAIBE and 0.384 for CRISPRon-BE; the corresponding CBE values are 0.845 and 0.191. The rank coefficients are also higher for crispAIBE: 0.481 versus 0.448 for ABE and 0.359 versus 0.228 for CBE. These comparisons concern a source absent from both models’ training data. Restricting the comparison to the benchmark’s designated high-frequency outcomes leaves crispAIBE with the higher *R*_2_ in seven of eight comparisons (Supplementary Table S1 and Note 2).

**Fig. 3.**
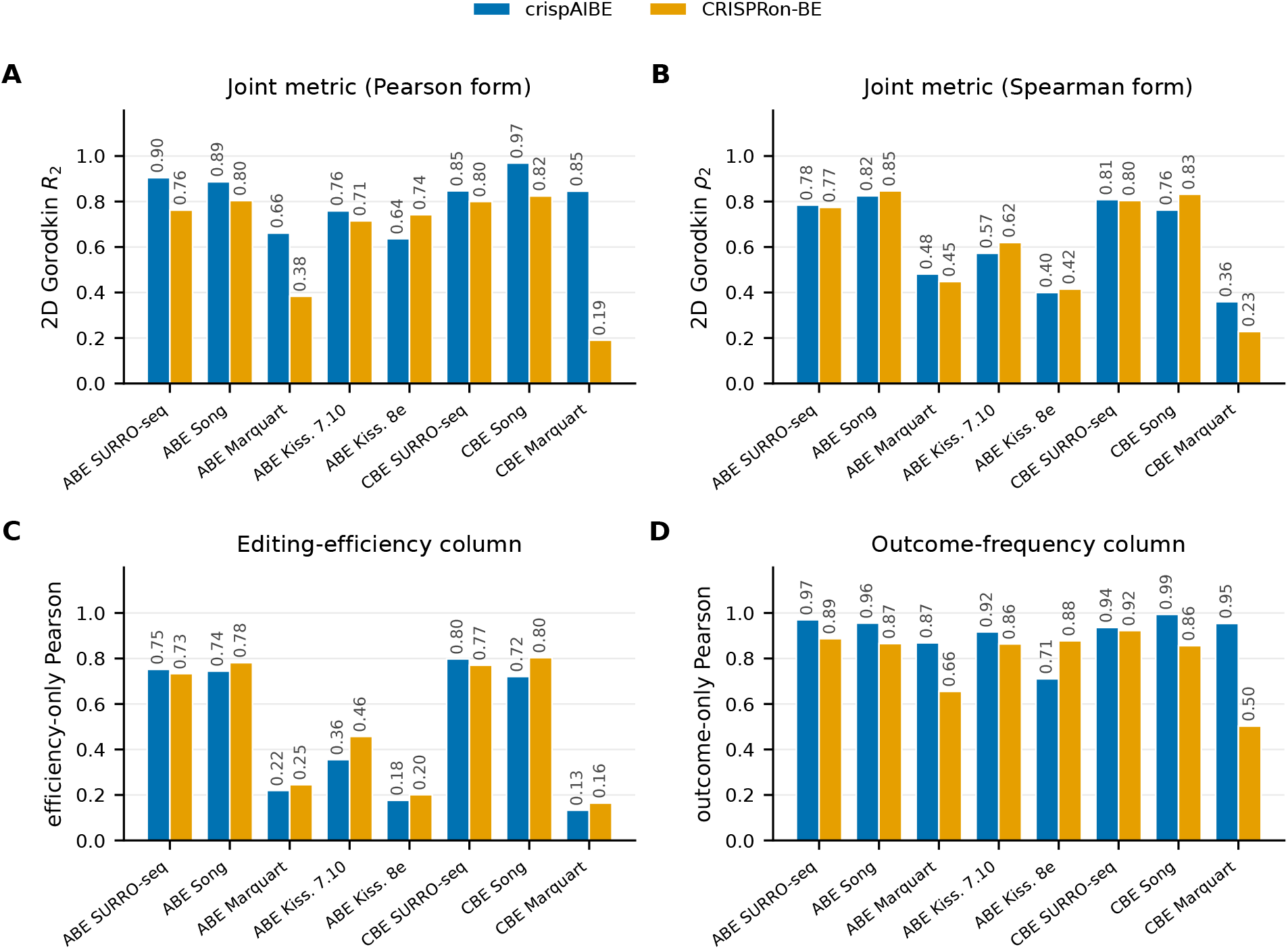
Comparison on the published benchmark test sets. **(A–B)** Two-column Pearson *R*_2_ and rank *ρ*_2_ coefficients. CRISPRon-BE values are the highest published dataset-label variant for each set and metric [30]. **(C)** Efficiency Pearson correlation after reducing to one row per guide. **(D)** Outcome-frequency Pearson over all outcome rows, including wild type. Panels C and D use different row weighting and are not an exact decomposition of panel A.

The individual targets help interpret these results (Fig. 3C–D). Across sets, crispAIBE’s outcome-frequency Pearson correlation ranges from 0.711 to 0.994. Guide-level efficiency Pearson is lower on Marquart (0.220 for ABE and 0.133 for CBE) and Kissling ABE8e (0.176), as it is for CRISPRon-BE (0.245, 0.164 and 0.201, respectively). CRISPRon-BE uses a 30-nt sequence context, calculated gRNA–DNA binding energy *ΔG*_*B*_, and predicted Cas9 efficiency [30]. Its authors report feature-removal ablations in their Supplementary Fig. 4 and Supplementary Data 3. The effects depend on the test set: with dataset-label settings held fixed, removing predicted Cas9 efficiency changes ABE *R*_2_ from 0.804 to 0.754 on Song, but from 0.715 to 0.755 on Kissling ABE7.10. These ablations describe feature contributions within CRISPRon-BE. Our benchmark efficiency model also uses engineered inputs (Section 2.2), and the cross-model comparison does not isolate the effects of input representation, architecture or assay differences. The source-balanced benchmark efficiency model attains higher Spearman correlation than source-specific gradient-boosting baselines in all nine eligible groups (Supplementary Table S2 and Note 2). The comparison uses that model’s modified-corpus test partition, a different evaluation population from the eight external benchmark comparisons above.

**Fig. 4.**
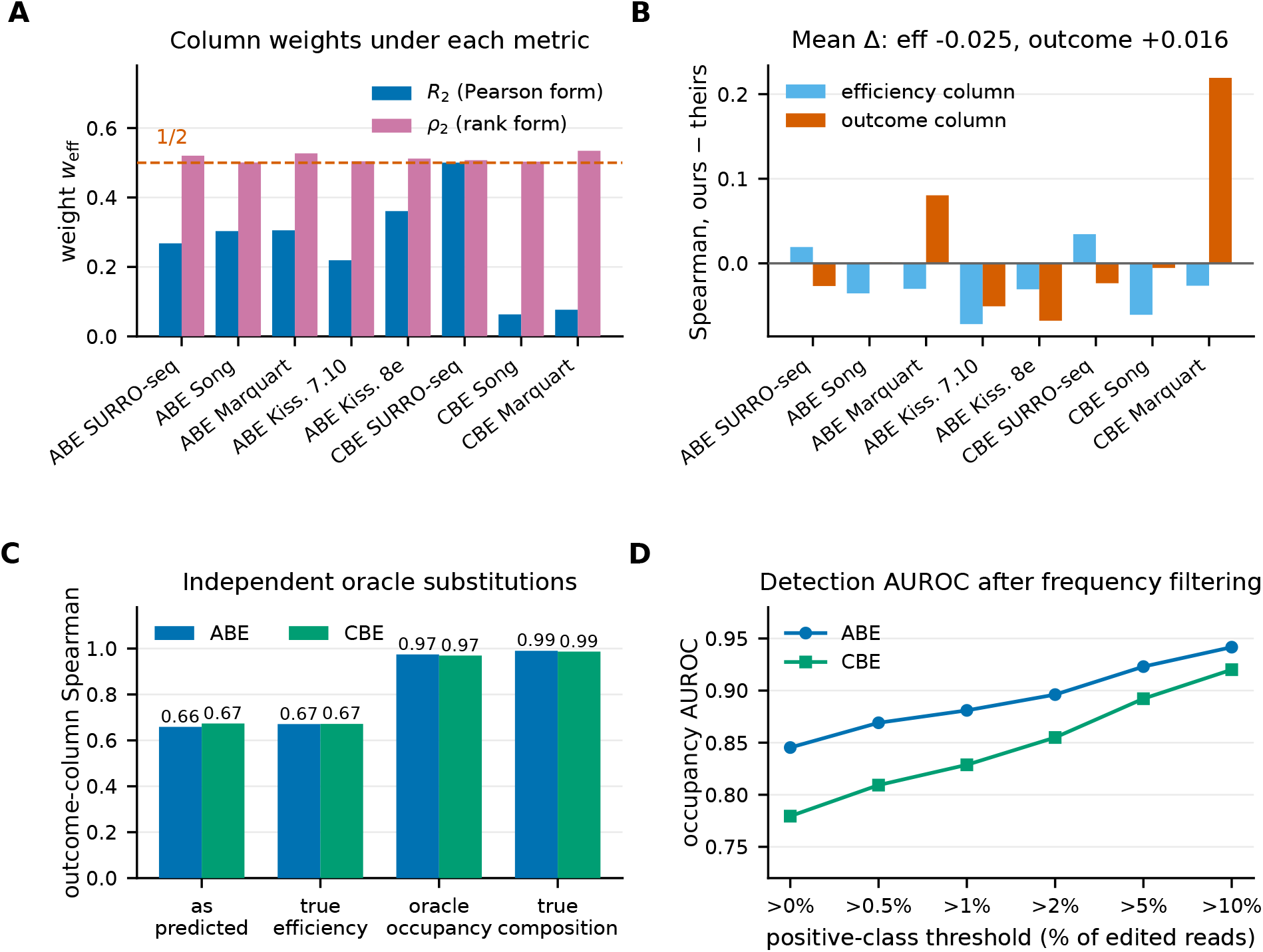
Interpretation of the joint benchmark statistics. **(A)** Coefficient assigned to the efficiency column under the Pearson and rank forms; the dashed line is 0.5. **(B)** Difference from CRISPRon-BE’s published per-target Spearman values. Efficiency is evaluated once per guide, so this panel does not exactly decompose the joint statistic. **(C)** Edited-outcome frequency Spearman after oracle substitutions, using the same composition model throughout and pooling the eight retained test sets by editor class. Each substitution starts from the original predictions: observed efficiency replaces efficiency alone; the observed-zero mask removes mass from unobserved edited alleles and renormalises; observed composition replaces composition alone. The latter two retain predicted efficiency. **(D)** AUROC for observed versus unobserved edited alleles after excluding positive alleles at or below the indicated conditional-frequency threshold. The retained classification population changes with the threshold.

### 3.4 Interpreting the two-column benchmark metrics

The two metric forms assign different coefficients to efficiency and outcome frequency. Writing 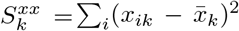 and defining 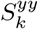analogously, Gorodkin’s coefficient is *R* = *Σ*_*k*_ *r*_*k*_ *w*_*k*_, where 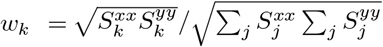 and *r*_*k*_ is the correlation in column *k*. These coefficients need not sum to one. Under the Pearson form, the efficiency coefficient ranges from 0.063 to 0.500 across the eight sets, while the outcome-frequency coefficient ranges from 0.491 to 0.932. Without ties, each within-column rank vector is a permutation of the same ranks and has the same sum of squared deviations. Each column coefficient is then one half, and the rank statistic equals the mean of the two column-wise Spearman correlations calculated over those outcome rows. Ties prevent exact equality in these data: the efficiency coefficient under ranking ranges from 0.501 to 0.535 (Fig. 4A). Thus *ρ*_2_ is close to this mean, whereas *R*_2_ gives outcome frequency more influence on most sets through its dependence on the raw-value scales and variances.

Outcome frequency itself combines different targets. Each guide contributes a wild-type row, whose frequency is the complement of editing efficiency, alongside its edited outcomes. High outcome-frequency correlation therefore does not isolate prediction of conditional bystander composition. Efficiency is also repeated once per outcome in the joint metric, so guides with more enumerated outcomes receive more weight. The per-target benchmark comparisons in Fig. 4B are descriptive because the published efficiency correlations use one row per guide.

To examine the outcome-frequency ranking, we substituted observed quantities for parts of the prediction while retaining the same edited-outcome rows (Fig. 4C). Replacing predicted efficiency with observed efficiency changes the ranking less than replacing the conditional composition. Setting predicted mass to zero for outcomes unobserved in the assay and renormalising raises outcome-frequency Spearman from 0.659 to 0.974 for ABE and from 0.673 to 0.969 for CBE. Using the full observed composition gives 0.990 and 0.987, respectively. These oracle substitutions use observed outcomes to diagnose sensitivity to the detected allele set. Their correlations describe diagnostic comparisons, not predictive performance or a lower bound on achievable error.

Among edited outcomes, the composition model distinguishes observed from unobserved alleles with AUROC 0.845 for ABE and 0.779 for CBE. Restricting the positive class to alleles above 10% conditional frequency raises these values to 0.942 and 0.920 (Fig. 4D). This restricted task distinguishes abundant observed alleles from unobserved alleles. Read depth is also associated with the observed labels: the zero-frequency fraction is 70.0% in the shallowest depth quintile (median 190 reads) and 47.2% in the deepest (median 5,407 reads). Detection AUROC decreases from 0.858 to 0.766 across these groups. Source, editor and outcome-set differences may confound these pooled associations.

Under a binomial sampling model, an allele with true frequency 0.002 has probability (1− 0.002)^1040^ ≃0.125 of yielding no reads at the median depth of 1,040 across distinct source–editor–guide combinations. This calculation illustrates how finite read depth can yield an observed zero for an allele with nonzero frequency. The pooled data do not separate this sampling contribution from model error or differences among assays.

### 3.5 Output-head and backbone ablations

With the remaining architecture fixed, the output heads attain Pearson correlations of 0.674 (point-MSE), 0.668 (Gaussian) and 0.667 (Dirichlet). With the Dirichlet output fixed, Pearson correlation is 0.667 for the transformer–CNN backbone, 0.663 for the transformer-only backbone and 0.655 for the CNN-only backbone, a range of 0.012. The Dirichlet parameterisation supplies a distribution supported on valid outcome proportions, with point accuracy close to the regression and Gaussian alternatives in these single-run comparisons. Supplementary Note 1 and Fig. S1A–B give the complete ablation results and calibration protocols.

### 3.6 Coverage depends on calibration population and editor stratification

Held-out calibration produced efficiency and composition coverage close to their nominal targets across all six evaluated levels (Table 3). Reusing the validation fold gives lower efficiency coverage at all six levels. That fold was involved in early stopping, and its source composition also differs from the test set because it contains no Marquart observations. The comparison therefore changes both predictor-selection independence and the calibration population. It does not isolate either effect.

**Table 3.**
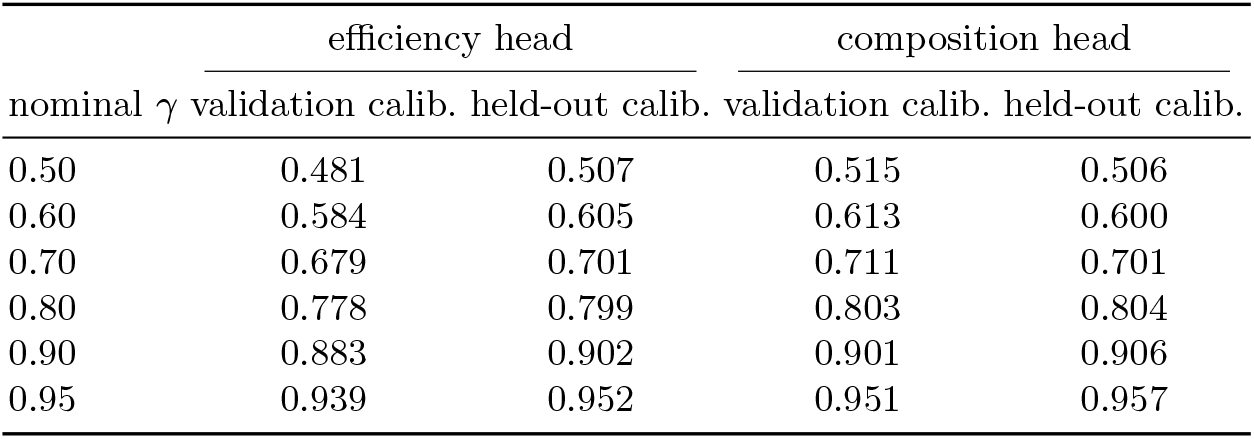
Empirical coverage of the split-conformal region against nominal *γ*. “Validation calibration” reuses the fold used for early stopping; “held-out calibration” uses a disjoint half of the test split reserved from model selection. Composition thresholds are fitted separately for each represented set size.

On the same efficiency checkpoint, we compared a model-based HDR, a pooled split-conformal region and an editor-stratified Mondrian region (Fig. S1C). At nominal 0.90, pooled empirical coverages are 0.887, 0.902 and 0.901, respectively. Mean region sizes are 0.384 for the model-based construction and 0.401 for the pooled conformal construction. The editor-level differences are larger: ABEmax coverage is 0.676 for the model-based region and 0.683 for pooled calibration, compared with 0.904 for Mondrian calibration. In this evaluation, editor-stratified calibration improved ABEmax coverage while maintaining near-nominal overall coverage.

The calibration source also matters. Under validation-fold calibration, the 306 ABEmax calibration observations exclude Marquart, although that source supplies 1,482 of 1,861 ABEmax test observations. Under this source mismatch, editor-specific calibration increased coverage from 0.662 to 0.690, remaining below the nominal 0.90. Coverage reached 0.904 when calibration and evaluation used the same mixture of screens.

### 3.7 Region area is associated with prediction error

Population-level coverage and guide-level error discrimination are distinct properties. We assessed the association between the efficiency model’s per-guide uncertainty and observed absolute error (Fig. 5). Regions in this section are calibrated on the validation fold, so that every guide in the test split carries an area and the ranking uses the full test split. These absolute areas are not directly comparable with the held-out calibration results in Section 3.6.

**Fig. 5.**
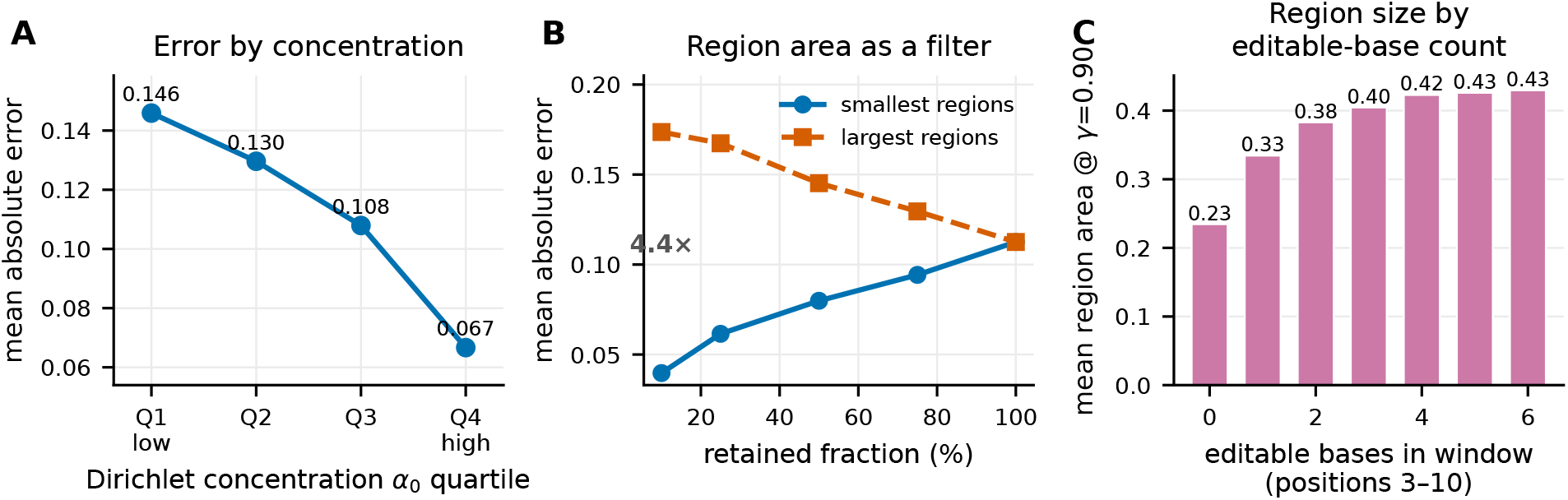
Region area and Dirichlet concentration as per-guide reliability signals. **(A)** Mean absolute error stratified by total Dirichlet concentration *α*_0_ quartile, decreasing monotonically from the least- to the most-concentrated quartile. **(B)** MAE for the smallest-versus largest-area fraction of predictions across retention levels; the annotation gives the ratio at 10% retention. **(C)** Mean conformal region area stratified by the number of editable bases in the positions 3–10 activity window.

Region area spans a factor of 3.8 between the fifth and ninety-fifth percentile of guides. It is positively associated with absolute error (Spearman *ρ* = 0.44) and total Dirichlet concentration *α*_0_ is negatively associated with it (*ρ* = −0.34). Across *α*_0_ quartiles, mean absolute error decreases monotonically: 0.146→ 0.130→ 0.108 → 0.067, a two-fold difference between the lowest and highest concentration quartiles (Fig. 5A). When predictions are ranked by region area, the smallest-area decile attains MAE 0.040 against 0.174 for the largest-area decile, a 4.4-fold MAE difference; the corresponding figures are 0.062 vs 0.167 at the quartile level and 0.080 vs 0.145 at the median (Fig. 5B). We report MAE rather than rank correlation under filtering because retaining a subset also changes the range of observed efficiencies and can affect Spearman correlation.

The pooled MAE comparison includes substantial variation in editing efficiency. The smallest-area decile has a mean observed edited fraction of 0.045 against 0.457 for the largest-area decile, so the filter preferentially retains low-efficiency guides, where absolute error also depends on the proximity of predictions and observations to the boundary. Ranking instead by the distance of the predicted efficiency from the nearest boundary, a ranking based only on the predicted mean, gives a 4.2-fold gap, and ranking at random gives 1.0. The pooled MAE difference is therefore largely reproduced by this point-prediction heuristic. Within quintiles of predicted efficiency, which reduce variation in boundary proximity, ranking by region area still separates error in every stratum. The arithmetic mean of the five ratios comparing the largest-area with the smallest-area decile is 1.9; the same within-quintile comparison using boundary distance gives 1.7. The boundary-distance result shows that efficiency-related variation persists within quintiles. These retrospective comparisons quantify error separation but do not establish an independent guide-selection benefit of region area beyond boundary distance.

In the fitted efficiency model, mean region area increases from 0.23 for guides with no editable bases in the represented window to 0.43 for guides with six (Fig. 5C). Window GC content shows little association with region area.

### 3.8 Uncertainty under cell-line transfer

We evaluated transfer to U2OS by excluding this cell line from training and validation and evaluating all U2OS records (Fig. 6). This experiment tests accuracy, coverage and the association between region area and error in a cell context absent from training. Of 14,593 U2OS test protospacers, 12,769 (87.5%) occur in training under another cell context. The evaluation therefore includes sequence overlap across cell contexts.

**Fig. 6.**
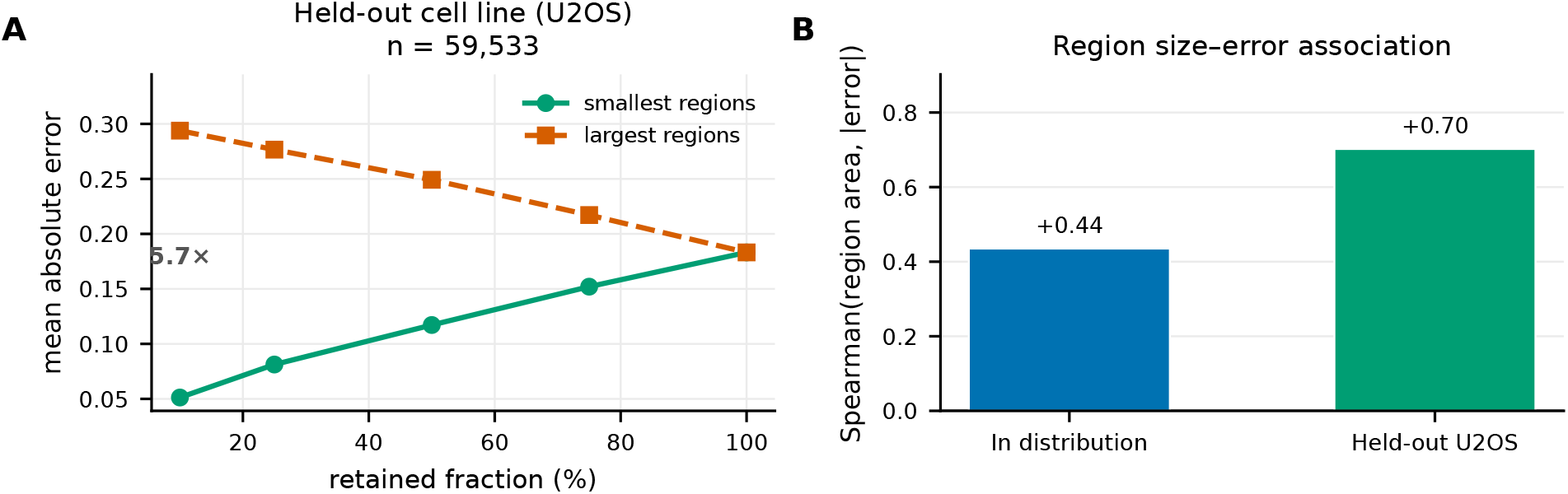
Uncertainty–error associations for transfer to U2OS. **(A)** With U2OS held out, the smallest- and largest-area deciles differ 5.7-fold in MAE; this pooled comparison is not conditioned on predicted efficiency. **(B)** Spearman correlation between region area and absolute error in distribution and for the U2OS holdout. The transfer results describe this cell-line holdout with sequence overlap across cell contexts.

#### Accuracy and error discrimination

With training restricted to HEK293T and mES, efficiency Spearman correlation is 0.38. Spearman correlation between region area and absolute error is 0.70, compared with 0.44 in distribution. The smallest-area decile has MAE 0.051 and the largest-area decile MAE 0.294, a 5.7-fold difference. This pooled comparison is unadjusted for predicted efficiency and is therefore subject to potential boundary-related confounding.

#### Coverage and recalibration

With a threshold calibrated on the HEK293T/mES validation fold, coverage on U2OS was 0.710, below the nominal 0.90. This analysis uses validation-fold calibration rather than the disjoint test-half calibration used for the main coverage estimate of 0.902 (Section 3.6).

We then recalibrated within U2OS using 300 random samples at each of five budgets (10, 25, 50, 100 and 250 distinct protospacers). Each sample uses one randomly selected record per protospacer; all records from those protospacers are removed from evaluation. With 100 calibration protospacers, median record-weighted coverage is 0.918 (10th–90th percentiles 0.879–0.951). At 250, the median is 0.918 (0.890–0.934). Local outcome measurements brought median coverage close to the nominal level under this sampling protocol. The median was similar at these two budgets, while the central 80% range across calibration samples narrowed at 250 protospacers. Recalibration leaves the point predictions unchanged; the calibration and evaluation units have different record weighting.

## 4 Discussion and Conclusions

Overall editing efficiency does not specify how much of a desired allele a guide will produce: guides with comparable activity can distribute their edited products differently among intended and bystander outcomes. crispAIBE addresses these two aspects of base editing through separately trained probabilistic models of efficiency and conditional allele composition. The framework extends the fixed outcome categories used in crispAIPE [21] to guide-dependent combinations of edited positions. Representing allele identity as an input allows the same composition network to score different outcome sets, while the Dirichlet distribution models their proportions jointly under the constraint that they sum to one.

This representation separates the expected composition from its predictive dispersion. The Dirichlet density provides a geometry for the predictive region, and held-out observations set its calibrated threshold. Conformal prediction provides the statistical basis for this calibration [2,25], including for compositional targets [1]. For guide-dependent allele sets, the number of components also matters: pooling density scores across dimensions can give approximately nominal overall coverage while under-covering small sets. Calibration by set size addresses this mismatch and gives near-nominal empirical coverage for sets of two to eight outcomes, with singleton regions retaining their sole composition (Supplementary Table S3). The single-run output-head and backbone comparisons show similar point correlations across the tested configurations (Supplementary Fig. S1A–B and Note 1). The reason for using the Dirichlet output is its representation of valid compositions and their dispersion, alongside the observed predictive accuracy.

The external comparisons show that this probabilistic formulation also gives competitive point predictions. crispAIBE ranked first in seven of eight comparisons under the two-column Pearson statistic and first or second in seven under the rank statistic. The comparison with CRISPRon-BE restricted to outcomes with observed frequency above 5% of all reads showed a similar pattern under the Pearson statistic (Supplementary Table S1 and Note 2). For efficiency, comparisons with source-specific gradient-boosting baselines on the separate modified-corpus test partition also favoured crispAIBE (Supplementary Table S2 and Note 2). These comparisons support the predictive performance of the fitted models across the evaluated sources, although differences in architecture and inputs prevent attributing the source-baseline result specifically to training across sources.

The metric decomposition clarifies which aspects of prediction these rankings assess. The Pearson and rank forms weight efficiency and outcome frequency differently, and repeating efficiency once per outcome gives more weight to guides with larger outcome sets. Outcome frequency also includes the wild-type fraction, which is determined by efficiency. A strong combined score consequently need not imply equally strong prediction of the distribution among edited alleles. Reporting the separate targets connects the benchmark to the biological distinction between inducing an edit and producing a particular edited product. The oracle substitutions further show that outcome-frequency ranking is sensitive to which alleles are detected. At finite read depth, an unobserved allele can have nonzero underlying frequency; the detected outcome set is therefore part of the measurement against which a prediction is judged.

Predictive coverage and error discrimination provide complementary information. Coverage describes performance across an evaluation population, whereas the association between region area and error tests whether larger regions identify less accurate predictions. Region area separated errors within predicted-efficiency quintiles, with a mean decile MAE ratio of 1.9, compared with 1.7 for distance from an efficiency boundary. The latter also reproduced most of the pooled separation. These retrospective results support interpreting region area alongside predicted activity: a narrow region around low efficiency describes a different experimental prospect from a narrow region around high efficiency. They do not establish an independent guide-selection benefit beyond the predicted-mean baseline.

The ABEmax and U2OS analyses show why the calibration population is part of this interpretation. Editor-specific calibration gave ABEmax coverage of 0.904 when calibration and evaluation drew on the same screens (Supplementary Fig. S1C), but did not reach nominal coverage when calibration excluded the laboratory supplying most test observations. Stratifying by editor alone therefore did not account for the differences between those screen populations. In the U2OS cell-context holdout, which includes sequence overlap with training, region area remained positively associated with error even as coverage fell to 0.710. Calibration on 100 U2OS protospacers brought median empirical coverage to 0.918 at nominal 0.90 without changing the predicted means. Local calibration therefore adjusted the extent of the predictive regions while leaving the model’s point accuracy unchanged. The positive uncertainty–error association and the initial undercoverage also show that error discrimination alone does not establish calibration in a new context.

The represented outcomes and sampling design define the scope of these conclusions. The main composition accuracy and coverage concern allele sets constructed from measured outcomes; sequence enumeration defines a different outcome space. The eight-position encoding captures combinations of edited positions while merging substitution identities and does not model indels explicitly. The reported coverage is also empirical and record-weighted: repeated protospacers across contexts mean that the exchangeable-observation guarantee does not directly establish coverage for this clustered evaluation. These distinctions matter when comparing screens or applying the framework to a different representation of editing products.

crispAIBE brings the distribution of edited products and the dispersion of their predictions into the same compositional framework. Its contribution is the combination of guide-specific allele modelling, competitive outcome prediction and calibration that accounts for the represented outcome set. Together, these provide a basis for assessing expected editing outcomes and predictive uncertainty in a specified experimental context.

## Acknowledgements

Furkan Özden acknowledges support through a Google DeepMind Scholarship.

## Funding

This work was supported by Google DeepMind.

## Disclosure of Interests

The authors have no competing interests to declare that are relevant to the content of this article.

## Supplementary material

The supplement comprises Supplementary Figure S1 (output-head, backbone and region-construction ablations), Supplementary Tables S1–S3 (the high-frequency benchmark, source-specific efficiency baselines and composition coverage by outcome-set size), and Supplementary Notes 1 and 2 (ablation protocols and additional benchmark comparisons).

## Data and code availability

The primary datasets were obtained from public releases. The BE-Hive screen is the figshare release of Arbab et al. [4] (articles 10673816 and 10678097). The multi-dataset compilation and partition-6 files come from the software and data release accompanying Sun et al. [30], archived at https://doi.org/10.5281/zenodo.16981783. Published comparator values come from their Supplementary Data 3. The trained models and scripts for corpus construction, model training, calibration, benchmark evaluation and figure generation are included in the accompanying reproducibility archive, together with prediction dumps and evaluation outputs. These resources are available at https://github.com/furkanozdenn/crispAIBE. The archive manifest maps results to configurations, checkpoints and source-data hashes. Primary screen data are obtained from the cited releases rather than redistributed in the archive.

## Notes

### Competing Interest Statement

The authors have declared no competing interest.

