## Supplementary Material for "Probabilistic Modelling of Base Editing Efficiency and Bystander Outcome Composition"

### Supplementary Figures

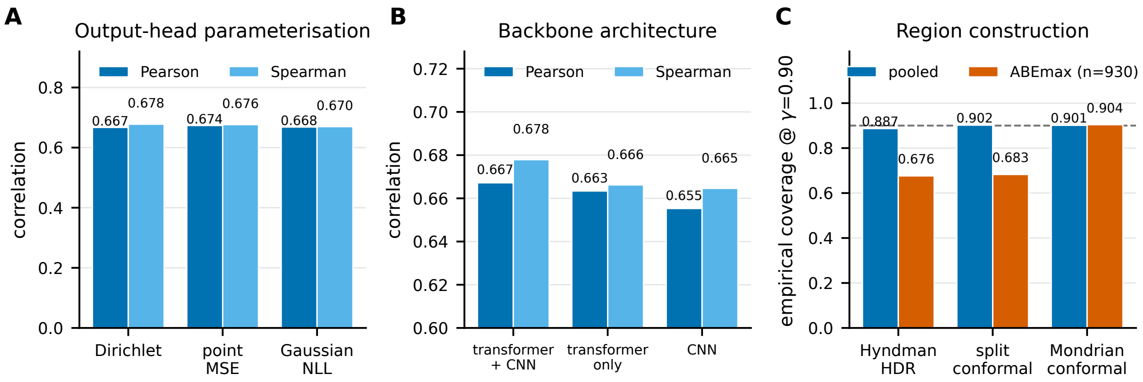


Figure S1: Efficiency-model ablations and region-construction comparisons. Prediction regions target coverage $\gamma = 0.90$. **(A)** Point-prediction accuracy for Dirichlet, point-MSE and Gaussian negative-log-likelihood heads. **(B)** Transformer + CNN, transformer-only and CNN-only backbones with a fixed Dirichlet head. **(C)** Pooled and ABEmax coverage for model-based HDRs, pooled split-conformal density level sets and editor-stratified Mondrian level sets. ABEmax coverage is 0.676, 0.683 and 0.904, respectively. Panels A and B use separately trained ablation models evaluated on the full test split; their conformal thresholds use the validation fold. Panel C uses the main efficiency checkpoint, with disjoint calibration and evaluation halves of the test set assigned by protospacer. Only the conformal constructions use the calibration half; the model-based HDR uses the predicted distribution directly. See Supplementary Note 1.

### Supplementary Tables

Table S1: Two-dimensional Pearson coefficient $R_2$ for outcomes with observed frequency $> 5\%$ of all reads, including wild-type outcomes meeting this threshold. These are the comparisons marked with an asterisk by CRISPRon-BE. CRISPRon-BE values are the best-performing dataset-label variant per test set in Supplementary Data 3 of Sun et al. (*Nature Communications* **16**, 9821, 2025; `https://doi.org/10.1038/s41467-025-65200-5`). The published test-set label "Kissling ABE7.10" is retained here; the corresponding model metadata uses ABEmax. $\Delta$ is crispAIBE minus CRISPRon-BE, calculated before rounding.

| Editor | Test set | crispAIBE | CRISPRon-BE | $\Delta$ |
|---|---|---|---|---|
| ABE | SURRO-seq | **0.890** | 0.744 | +0.147 |
| ABE | Song | **0.856** | 0.733 | +0.123 |
| ABE | Marquart | **0.546** | 0.255 | +0.291 |
| ABE | Kissling ABE7.10 | **0.770** | 0.626 | +0.143 |
| ABE | Kissling ABE8e | 0.444 | **0.681** | −0.237 |
| CBE | SURRO-seq | **0.837** | 0.796 | +0.042 |
| CBE | Song | **0.973** | 0.716 | +0.257 |
| CBE | Marquart | **0.820** | 0.146 | +0.673 |

Table S2: Spearman correlation for the efficiency model trained across sources and source-specific gradient-boosted trees using one-hot encoded 20-nt protospacers. The crispAIBE model is the benchmark-excluded `crispron_holdout_bal05` checkpoint (epoch 8). Both models are evaluated on the full test partition of its modified corpus, rather than only the eight external benchmark sets. Each baseline is fitted to all labelled training records in its source × stored editor-category group; editor variants and cell types are pooled within each group. All nine groups with at least 200 training records and 50 test records are shown. Counts refer to records, not distinct protospacers. The CtoGA library is stored under the historical category `CGBE`; this label does not identify the molecular editor class. $\Delta$ is crispAIBE minus the baseline, calculated before rounding.

| Source | Category | Train records | Test records | crispAIBE | Baseline | $\Delta$ |
|---|---|---|---|---|---|---|
| Arbab | ABE | 86,350 | 20,876 | **0.614** | 0.474 | +0.140 |
| Arbab | CBE | 286,427 | 68,853 | **0.660** | 0.293 | +0.368 |
| Arbab | CtoGA library | 9,466 | 562 | **0.696** | 0.461 | +0.235 |
| Kissling ABE8e | ABE | 723 | 239 | **0.161** | 0.011 | +0.150 |
| Kissling ABEmax | ABE | 687 | 224 | **0.499** | 0.455 | +0.044 |
| Song | ABE | 4,479 | 696 | **0.734** | 0.568 | +0.166 |
| Song | CBE | 3,560 | 689 | **0.732** | 0.599 | +0.133 |
| SURRO-seq | ABE | 3,974 | 2,564 | **0.766** | 0.656 | +0.110 |
| SURRO-seq | CBE | 5,856 | 2,598 | **0.801** | 0.764 | +0.037 |

Table S3: Composition-region coverage at nominal 0.90 by represented allele-set size $K$. Both methods use the same disjoint test-set halves, assigned by protospacer. Pooled density uses one negative-log-density threshold across $K$; the main-text method calibrates within each $K$ and always retains the singleton simplex for $K = 1$. Entries are the fraction of evaluation records whose observed composition lies in the region.

| $K$ | Calibration records | Evaluation records | Pooled density | By set size |
|---|---|---|---|---|
| 1 | 102 | 117 | 0.000 | 1.000 |
| 2 | 623 | 629 | 0.523 | 0.905 |
| 3 | 721 | 673 | 0.761 | 0.896 |
| 4 | 1,239 | 1,335 | 0.881 | 0.910 |
| 5 | 823 | 968 | 0.952 | 0.897 |
| 6 | 776 | 863 | 0.980 | 0.911 |
| 7 | 664 | 752 | 0.987 | 0.895 |
| 8 | 2,807 | 2,930 | 0.995 | 0.909 |
| All | 7,755 | 8,267 | 0.900 | 0.906 |

### Supplementary Note 1: Output-head and backbone ablations

Each configuration was trained once; variability across training runs was not estimated.

**Output head.** With the backbone fixed (Fig. S1A), Pearson and Spearman correlations are 0.674 and 0.676 for point-MSE, 0.668 and 0.670 for Gaussian negative log likelihood, and 0.667 and 0.678 for the Dirichlet head. Gaussian and Dirichlet region coverage is 0.883 and 0.884, respectively; no region was constructed for point-MSE. Dirichlet regions use negative-log-density scores; Gaussian intervals use standardised absolute residuals $|y - \mu|/\sigma$, where $\mu$ and $\sigma$ are the predicted mean and standard deviation.

The output-head and backbone runs used the validation fold for both model selection and calibration; their coverage estimates are empirical diagnostics. The region-construction comparison in Fig. S1C instead follows the main-text split-conformal analysis, with disjoint test-set halves assigned by protospacer.

**Backbone.** With a fixed Dirichlet head, Pearson correlation is 0.667 for transformer + CNN, 0.663 for transformer-only and 0.655 for CNN-only; coverage ranges from 0.880 to 0.884 (Fig. S1B).

### Supplementary Note 2: Additional benchmark comparisons

**High-frequency outcomes.** On outcomes with observed frequency above 5% (Table S1), crispAIBE has higher $R_2$ in seven of eight comparisons; its $R_2$ is lower for Kissling ABE8e by 0.237. The main-text benchmark exclusions apply.

**Source-specific baseline.** crispAIBE has higher Spearman correlation in all nine eligible groups (Table S2). Baselines used `HistGradientBoostingRegressor` (scikit-learn 1.6.1) with at most 400 iterations, learning rate 0.06, seed 0 and remaining defaults. Because architecture and inputs also differ, this comparison does not isolate the effect of training across sources.
